# Single laboratory evaluation of the (Q20+) nanopore sequencing kit for bacterial outbreak investigations

**DOI:** 10.1101/2024.07.17.603985

**Authors:** Maria Hoffmann, Jay Hee Jang, Sandra Tallent, Narjol Gonzalez-Escalona

## Abstract

This study aimed to evaluate the potential of Oxford Nanopore Technologies (ONT) GridION with Q20+ chemistry as a rapid and accurate method for identifying and clustering foodborne pathogens. The study focuses on assessing whether ONT Q20+ technology could offer near real-time pathogen identification, including SNP differences, serotypes, and antimicrobial resistance genes, to overcome the drawbacks of existing methodologies. This pilot study evaluated different combinations of two DNA extraction methods (Maxwell RSC Cultured Cell DNA kit, and Monarch high molecular weight extraction kits) and two ONT library preparation protocols (ligation and the rapid barcoding sequencing kit) using five well-characterized strains representing diverse foodborne pathogens. The results showed that any combination of extraction and sequencing kits produced high-quality closed bacterial genomes. However, there were variations in assembly length and genome completeness based on different combinations of methods, indicating the need for further optimization. *in silico* analyses demonstrated that the Q20+ nanopore sequencing chemistry accurately identified species, genotyped, and detected virulence factors comparable to Illumina sequencing. Phylogenomic clustering methods showed that ONT assemblies clustered with reference genomes, although some indels and SNP differences were observed. There were also differences on SNP accuracy among the different species. The observed SNP differences were likely due to sequencing and analysis processes rather than genetic variations in the sampled bacteria. The study also compared a change in the basecaller model with the previous model (SUP 4Khz 260 bps) and found no significant difference in accuracy (SUP 5Khz 400 bps). In conclusion, the evaluation of ONT Q20+ nanopore sequencing chemistry demonstrated its potential as an alternative for rapid and comprehensive bacterial genome analysis in outbreak investigations. However, further research, verification studies, and optimization efforts are needed to address the observed limitations to adopt and fully realize the impact of nanopore sequencing on public health outcomes and more efficient responses to foodborne disease threats.

## Introduction

Leafy greens are responsible for nearly half of the produce-related Shiga toxin-producing *Escherichia coli* (STEC) outbreaks in the United States and recent investigations have implicated agricultural water as a potential source. Currently, the Food and Drug Administration (FDA) outbreak detection protocols for STEC require extensive analysis time (2 – 4 weeks). During outbreak investigations finding the strain that matches the outbreak strain can take several days after isolating them from the potential food sources. This can be very crucial considering that fresh produce cannot be held for long, and during outbreak events and uncertainty, methods are needed to quickly identify unsafe products.

Currently, the FDA utilizes Illumina instruments for whole genome sequencing (WGS) analysis of isolated strains for surveillance [1, 2]. While short-read Illumina MiSeq sequencing technology is highly accurate, it struggles with assembling highly repetitive regions that can span several hundred base pairs [3–7]. As a result, the final assembled genome of an outbreak strain typically consists of numerous contigs (e.g., approximately 200 contigs for STECs) and often lacks critical information such as plasmid presence and the location of important genes, including antimicrobial resistance genes (AMR) [7–9]. An alternative approach involves using long reads from Oxford Nanopore Technologies (ONT), which can identify taxa while sequencing. This method has shown promise outbreak detection [5, 10], but its lower raw read accuracy previously made it unsuitable as a stand-alone technique for certain downstream analyses, such as SNP analysis during outbreak investigations [1, 2]. Accurate SNP variant detection is crucial in outbreak investigations, as some strains can differ by only a few SNPs (around 1-10) [1, 2]. However, in 2023, ONT released a new chemistry (Q20+), achieving ≥99% raw read accuracy and high sequencing yield [11–14], making it a potential candidate for improved and faster outbreak response.

Prior to implementation of ONT sequencing as an alternative as an enhanced and faster tool for outbreak response, it is crucial to ensure that the quality obtained is in par with the current gold standard, Illumina sequencing. To achieve this, a pilot study of ONT’s new chemistry including the V14 DNA library preparation kits and new R10.4.1 flow cell is necessary to select the optimal combination to perform a more rigorous single laboratory validation (SLV). According to the FDA’s Guidelines for the Validation of Analytical Methods Using Nucleic Acid Sequenced-Based Technologies [15] several important considerations must be taken into account when conducting a verification study for a new change in next generation sequencing (NGS) chemistry such as the Q20+ from ONT. These considerations include: 1) the use of a well characterized isolate panel for primary NGS/WGS data collection, 2) reproducibility (precision between runs), and 3) repeatability (precision within a run). In this pilot study, these requirements will be initially evaluated against a set of reference genomes based on parameters such as: a) average depth of genome coverage, b) mean read length, c) sequence length distribution, d) assembly length, e) number of contigs, f) N50, and g) percentage of recovered open reading frames (ORF) and core genes. Subsequently, in silico tests will be conducted to assess correct species identification (accuracy, specificity, and sensitivity), genotyping methods (serotyping, AMR, virulence, toxin, multi-locus sequence typing (MLST), and phylogenomic clustering methods measuring specificity and sensitivity for cluster assignment. A wgMLST approach using the known genome reference of the tested strains as a reference will be employed to determine specificity and sensitivity for calling variants (single nucleotide polymorphisms (SNPS) and/or indels), and the accuracy of the entire tree or specific splits leading to an outbreak lineage. The entire workflow is depicted in figure 1.

**Figure 1.**
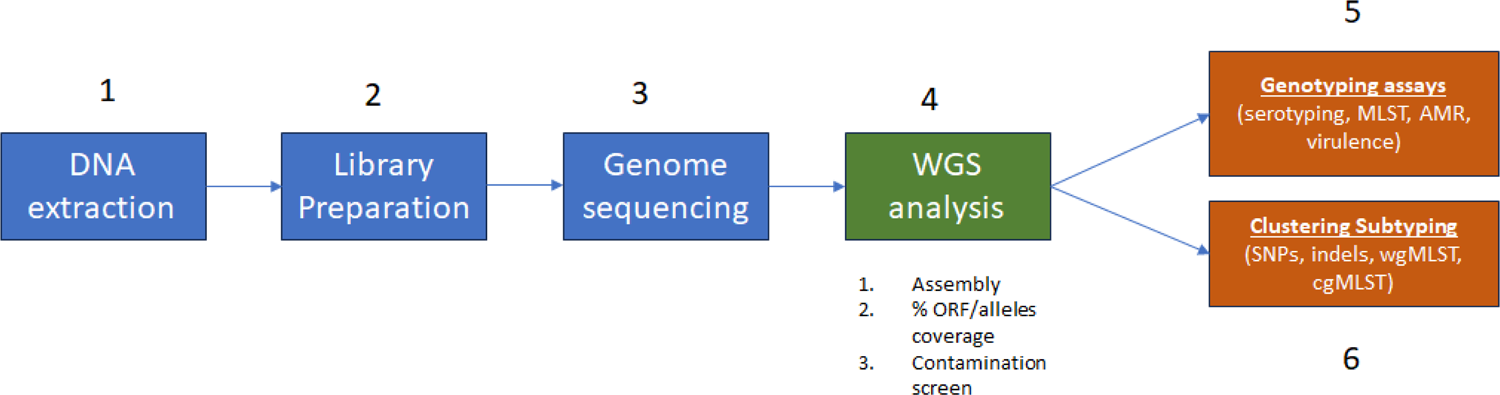
NGS Validation Workflow for Pure Bacterial Isolates according to Guidelines for the Validation of Analytical Methods for Nucleic Acid Sequence-Based Analysis of Food, Feed, Cosmetics and Veterinary Products from the FDA (https://www.fda.gov/food/laboratory-methods-food/foods-program-methods-validation-processes-and-guidelines).

ONT has two main DNA library preparation kits for whole genome sequencing of bacteria: 1) the Ligation sequencing kit (SQK-LSK114) and 2) the Rapid barcoding sequencing kit (SQK-RBK11). They differ in the preparation time and steps required to obtain a library ready for sequencing. The RBK kit is faster and less labor-intensive than the LSK kit, albeit producing lower sequencing output. The sequencing performance and output can be highly impacted by the quality of the DNA used for the sequencing. Therefore, we chose to also evaluate the performance of these two library preparation kits with DNA extracted with kits that produced DNA with different specifications: 1) faster and automatized Maxwell RSC Cultured Cell DNA kit (high DNA concentration output but sheared DNA) and 2) the manual Monarch high molecular weight (HMW) DNA extraction kit (DNA concentration high and with more integrity – longer fragments). For this pilot study we selected to sequence five well known and characterized foodborne bacterial pathogen strains belonging to five different species from our collection: *Salmonella enterica* subsp. *enterica*, *Vibrio parahemolyticus*, *Shigella sonnei*, *Escherichia coli*, and *Klebsiella pneumoniae* (Table 1). The DNA was extracted from overnight broth cultures with the two different kits and were sequenced using the two ONT library preparation kits separately, for a total of 4 runs as shown in Table 2. Therefore, in this pilot study we only conducted some assays that will test the first two requirements as per the FDA NGS validation guidelines: 1) the use of a well characterized isolate panel for primary NGS/WGS data collection, and 2) reproducibility (Precision between runs), with a later SLV that will test all three requirements with a more extended and inclusive foodborne bacterial strain panel.

**Table 1.**
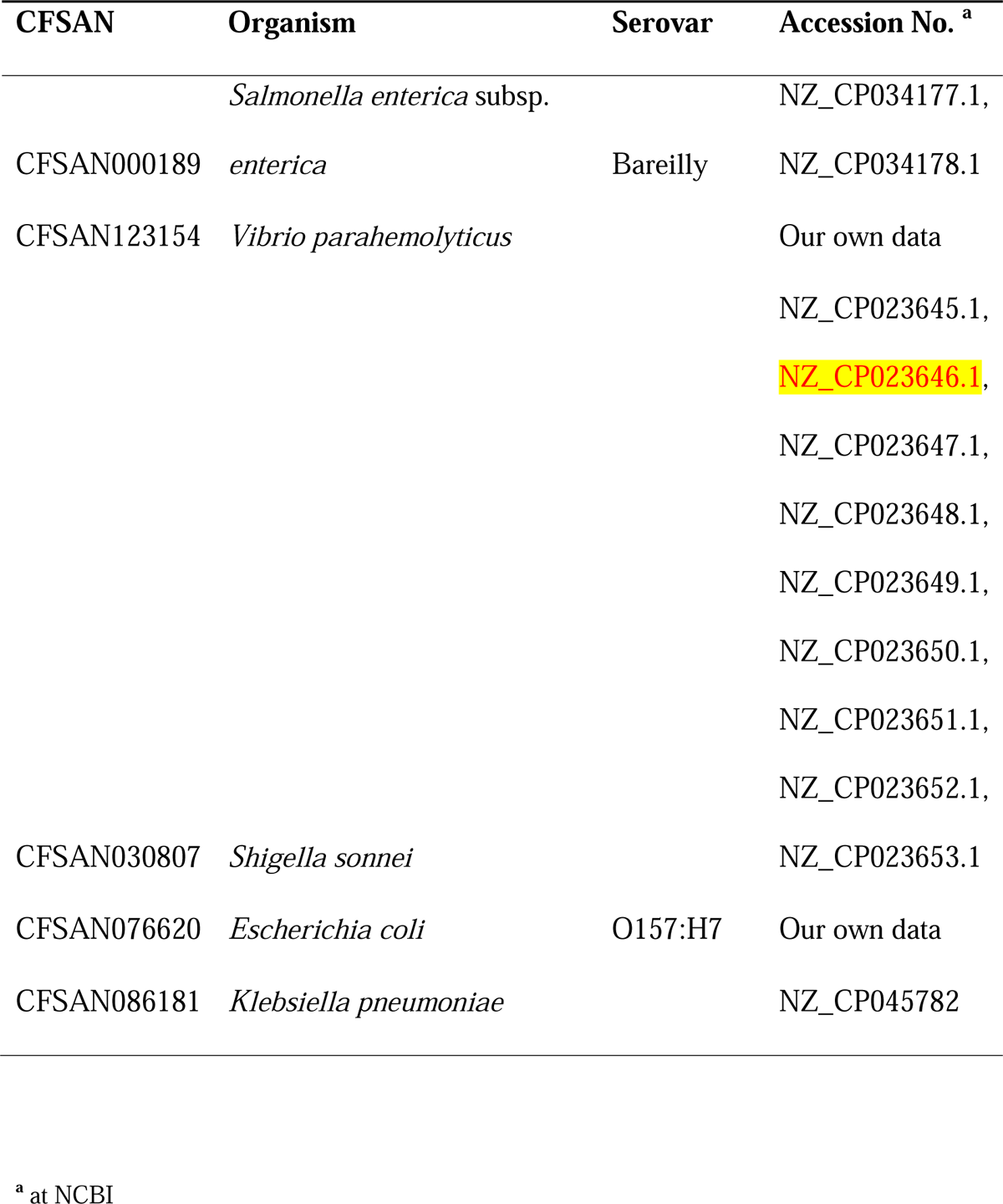
Verification strains set used in this single laboratory evaluation study.

| CFSAN | Organism | Serovar | Accession No. <sup>a</sup> |
| --- | --- | --- | --- |
|  | *Salmonella enterica* subsp. |  | NZ_CP034177.1, |
| CFSAN000189 | *enterica* | Bareilly | NZ_CP034178.1 |
| CFSAN123154 | *Vibrio parahaemolyticus* |  | Our own data |
|  |  |  | NZ_CP023645.1, |
|  |  |  | NZ_CP023646.1, |
|  |  |  | NZ_CP023647.1, |
|  |  |  | NZ_CP023648.1, |
|  |  |  | NZ_CP023649.1, |
|  |  |  | NZ_CP023650.1, |
|  |  |  | NZ_CP023651.1, |
|  |  |  | NZ_CP023652.1, |
| CFSAN030807 | *Shigella sonnei* |  | NZ_CP023653.1 |
| CFSAN076620 | *Escherichia coli* | O157:H7 | Our own data |
| CFSAN086181 | *Klebsiella pneumoniae* |  | NZ_CP045782 |

<sup>a</sup> at NCBI

**Table 2.** Initial testing of the Q20+ chemistry (RBK114 or LSK114 with R10.4.1 flow cells)

| Runs | Flowcell<br>No. | DNA<br>extraction<br>kit | Library prep<br>kit<sup>a</sup> | Basecalling<sup>b</sup> | Output | Read No. |
| --- | --- | --- | --- | --- | --- | --- |
| 1 | FAW43075 | Maxwell | SQK-LSK114 | SUP 260 bps | 10.65 Gb | 1.76 M |
| 2 | FAW25027 | Maxwell | SQK-RBK114.24 | SUP 260 bps | 4.3 Gb | 705.61 k |
| 3 | FAW42567 | NEB | SQK-LSK114 | SUP 260 bps | 8.83 Gb | 879.33 k |
| 4 | FAW18242 | NEB | SQK-RBK114.24 | SUP 260 bps | 3.28 Gb | 305.31 k |

Maxwell- Maxwell RSC Cultured Cell DNA kit, NEB- Monarch high molecular weight (HMW). <sup>a</sup>We used 50 ng per strain per flowcell in the case of the RBK kit and 400 ng for the LSK, as recommended by the manufacturer's protocols for each kit. <sup>b</sup>SUP- Super-accurate, <sup>a</sup>MinKNOW software version 22.12.5, basecall\_model - dna\_r10.4.1\_e8.2\_260bps\_sup@v3.5.2.

## MATERIALS AND METHODS

### Bacterial strains

The bacterial strains used in this study belonged to five different species associated with foodborne outbreak or cases collected by the Center for Food Safety and Applied nutrition (CFSAN) as part of our Genome Trakr surveillance program (Table 1). These five strains were selected to represent various species, antimicrobial resistance (AMR) genes, common contaminant, diverse %GC content (medium, and high), SNP recovery as part of the same outbreak, toxin gene detection, and diverse plasmid content. Bacterial strains were cultured in BD Difco™ brain heart infusion (BHI) broth (Thermo Fisher Scientific, Waltham, MA) and stored in BHI broth with 20% glycerol stocks at −80 C.

### DNA extraction, quality and quantify measurement

Genomic DNA from each strain was extracted using either the Maxwell RSC Cultured Cell DNA kit with a Maxwell RSC Instrument (Promega Corporation, Madison, WI) according to manufacturer’s instructions for gram-negative bacteria with additional RNase treatment or the Monarch HMW DNA Extraction kit (New England Biolabs, Ipswich, MA). DNA concentration was determined by Qubit 4 Fluorometer (Invitrogen, Carlsbad, CA) according to manufacturer’s instructions. The DNA integrity was determined using a Femto Pulse Systems (Agilent, Santa Clara, CA).

### Whole genome sequencing and assembly using long and reads

DNA extracted from each strain was sequenced using the GridION Sequencing Device Mk1 (Oxford Nanopore Technologies, Oxford, UK) in different conformations (Table 2). The sequencing libraries were prepared using different protocols: 1) Rapid Barcoding Kit 24 V14 (SQK-RBK114.24), or 2) Ligation Sequencing Kit v14 (SQK-LSK114) with Native Barcoding Kit (EXP-NBD114). The sequencing libraries were added to FLO-MIN114 (R10.4.1) flow cells, according to the manufacturer’s instructions for 48 hours (Oxford Nanopore Technologies). The sequencing run was monitored using the GridION’s live base called MinKNOW software versions (see Tables) using the super-accurate basecalling model (SUP). The sequenced reads that were < 2000 bp and quality scores of <10 were discarded for downstream analysis using Fitlong v0.2.1 (https://github.com/rrwick/Filtlong) with default parameters. The genomes for each strain were obtained by *de novo* assembly using the Flye program v2.9.2 [16], with the following parameters: --nano-hq, --read-error 0.03, and -i 4.

The short read whole genome sequences for these strains were generated by Illumina MiSeq sequencing with the MiSeq V3 kit using 2 x 250 bp paired-end chemistry, (Illumina, San Diego, CA) according to manufacturer’s instructions, at 100X coverage. The libraries were constructed using 100 ng of genomic DNA using the Illumina® DNA Prep (M) Tagmentation (Illumina, San Diego, CA), according to manufacturer’s instructions. Reads were trimmed with Trimmomatic v0.36 [17].

### Species identification

The assembled genome from every run was taxonomically classified by Kraken 2 [18] implemented in the GalaxyTrakr website [19].

### *in silico* MLST and serotyping

The initial analysis and identification for each strain was performed using an *in silico* MLST approach using Ridom SeqSphere+ v9.0.8 (Ridom, Münster, Germany) and Ridom’s MLST task template for each individual species as follows: for *Salmonella enterica* (https://enterobase.warwick.ac.uk/species/index/senterica), *Vibrio parahemolyticus* (https://pubmlst.org/organisms/vibrio-parahaemolyticus), *Shigella sonnei* (https://enterobase.warwick.ac.uk/species/index/ecoli), *Escherichia coli* (https://enterobase.warwick.ac.uk/species/index/ecoli) and *Klebsiella pneumoniae* (https://bigsdb.pasteur.fr/klebsiella/). (*Salmonella* and *E. coli*). Only two serotyping schemes were available for the 5 different species included in this pilot study (*Salmonella* and *E. coli*). For *Salmonella enterica* we used SeqSero2 v1.1.0 (http://www.denglab.info/SeqSero2) and for *E. coli* we used SerotypeFinder 2.0 (https://cge.food.dtu.dk/services/SerotypeFinder/) [20].

### *in silico* AMR gene detection

The AMR gene detection was performed using ResFinder v4.5.0 (http://genepi.food.dtu.dk/resfinder) [21].

### Phylogenetic relationship of the strains by wgMLST analysis

The phylogenetic relationship of the strains was assessed by a whole genome multilocus sequence typing (wgMLST) analysis using Ridom SeqSphere+ v9.0.8. A separate wgMLST was created for each individual species using the genome available at NCBI or obtained in house for that same strain as reference (Table 1). Genes that are repeated in more than one copy in any of the two genomes were removed from the analysis as failed genes. A task template was then created that contains both core and accessory genes for each reference strain for any future testing. Each individual locus (core or accessory genes) from the reference strain was assigned allele number 1. The assemblies for each individual strain closed genome in this study were queried against the task template for each individual species/strain. If the locus was found and was different from the reference genome or any other queried genome already in the database, a new number was assigned to that locus and so on. After eliminating any loci that were missing from the genome of any strain used in our analyses, we performed the wgMLST analysis. These remaining loci were considered the core genome shared by the analyzed strains. We used Nei’s DNA distance method [22] to calculate the matrix of genetic distance, considering only the number of same/different alleles in the core genes. A Neighbor-Joining (NJ) tree was built using pairwise ignoring missing values and the appropriate genetic distances after the wgMLST analysis. wgMLST uses the alleles number of each locus to determine the genetic distance and build the phylogenetic tree. The use of allele numbers reduces the influence of recombination in the dataset studied and allows for fast clustering determination of genomes.

### Nucleotide sequence accession numbers

The raw ONT data for each individual sample were deposited in GenBank under BioPro-ject accession number PRJNA1134675.

## RESULTS

### Preliminary testing of the Q20+ chemistry to establish which sequencing kit and DNA combination could be used for easily and completely close bacterial genomes for later validation efforts

We selected 5 strains from different species and GC content (CFSAN000189 - *Salmonella* Bareilly, CFSAN123154 - *Vibrio parahemolyticus*, CFSAN030807 - *Shigella sonnei*, CFSAN076620 - *E. coli* O157:H7, and CFSAN086181 - *Klebsiella pneumoniae*) for the initial ONT Q20+ chemistry single laboratory evaluation. As mentioned in materials and methods all sequencing runs were performed on a GridION (Ligation sequencing kit - SQK-LSK114 and Rapid barcoding sequencing kit SQK-RBK114.24, referred here as LSK114 or RBK114 moving forward) (Table 2). As shown in Table 2 each combination resulted in different total output and total read numbers. The highest output both in read numbers and sequence was achieved with the LSK114/Maxwell combination (10.65 Gb and 1.76 M reads). The lowest was obtained with the RBK114/NEB combination (3.28 Gb and 305k reads).

The higher N50 length reads per strain per run were observed on the runs using the DNA extracted with the NEB HMW kit with any of the two DNA library combination (Figure 1). The higher number of reads per strain per run were observed on the runs using the Maxwell/LSK114 combination. This explains partially why the RBK114/NEB run had the lowest output.

### Genome Sequencing QC

The data generated from the 4 runs was compared for assembly statistics: a) assembly length (bp), b) average depth of genome coverage, c) genome completeness, d) number of contigs, and e) % ORF genes recovered.

Table 3 shows the assembly length for each strain per run and the average depth coverage for each genome per run. The assembly length per strain per run showed a great variation. For the *Salmonella* strain the assembled genome length varied from 2 kb less to 5 kb more compared to its reference genome; for the *Vibrio parahemolyticus* from 5 kb to 38 kb more compared to the reference genome; for the *Shigella sonnei* from 17 kb less to 21 kb more compared to its reference genome; for the *E. coli* O157:H7 from 2 kb less to 22 kb more compared to its reference genome and for the *Klebsiella pneumoniae* the variation was less pronounced from 0.2 kb to 3.8 kb more compared to its reference genome. These variations do not appear to be linked to the sequencing depth, since for example *Klebsiella pneumoniae* showed the higher variation for the highest coverage (> 300X).

**Table 3.**
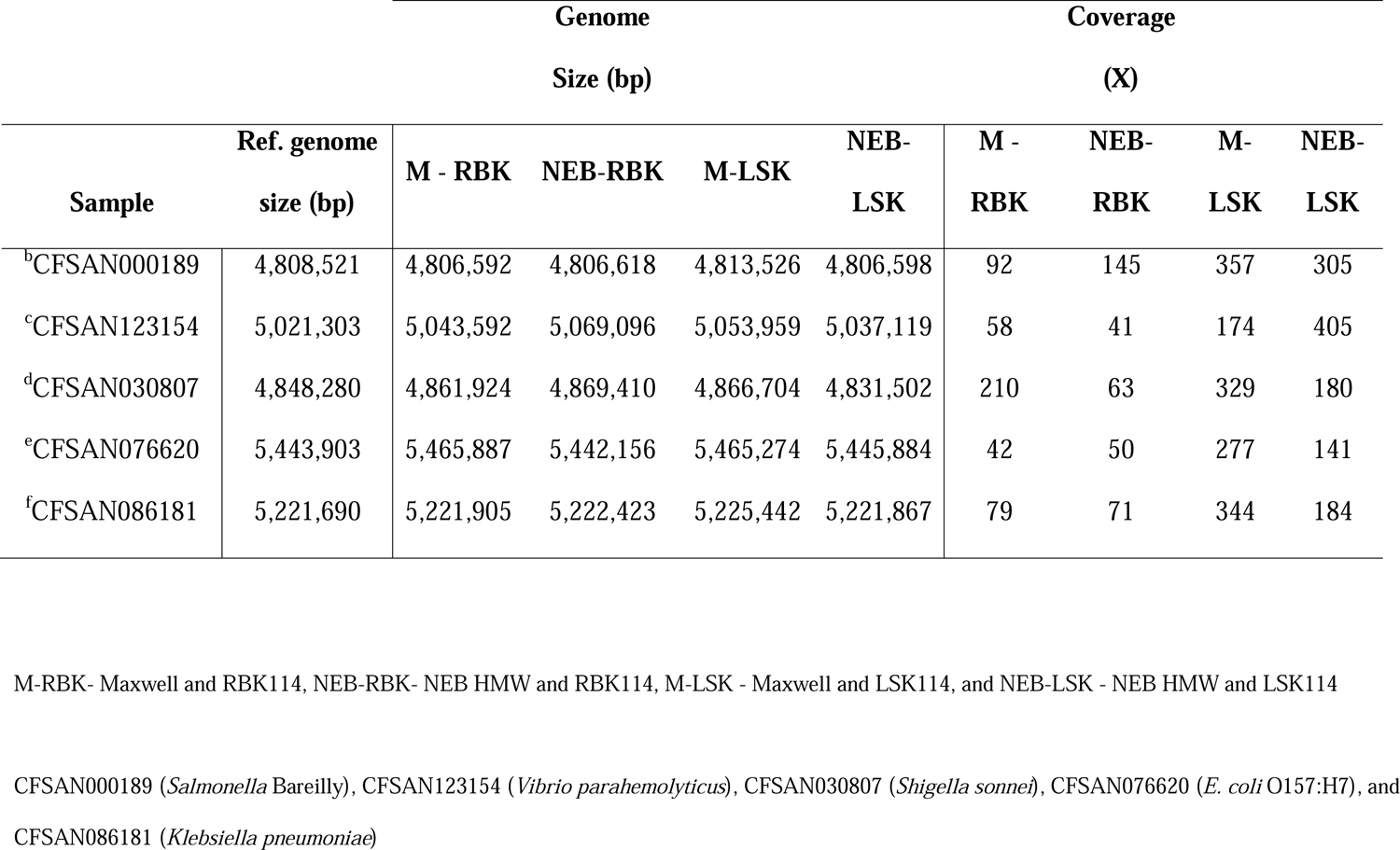
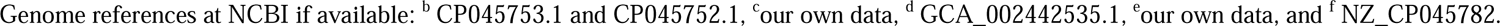
Assembly statistics showing genome size and coverage for each sample sequenced using any of the 4 combinations tested.

Regarding genome completeness, most genomes were complete except for CFSAN076620 (*E. coli* O157:H7, run RBK114-NEB and LSK114-NEB) and CFSAN086181 (*Klebsiella pneumoniae*, run RBK114-NEB) (Table 4). Most of the ORF genes (>99%) were recovered by ONT Q20+ in any of the combinations compared to the reference genome for each strain (Table 5). Based on the analyzed genome and sequencing combinations, there were still issues with indels and the recovery of some ORFs was not possible. The results were not identical and showed inconsistencies between runs.

**Table 4.**
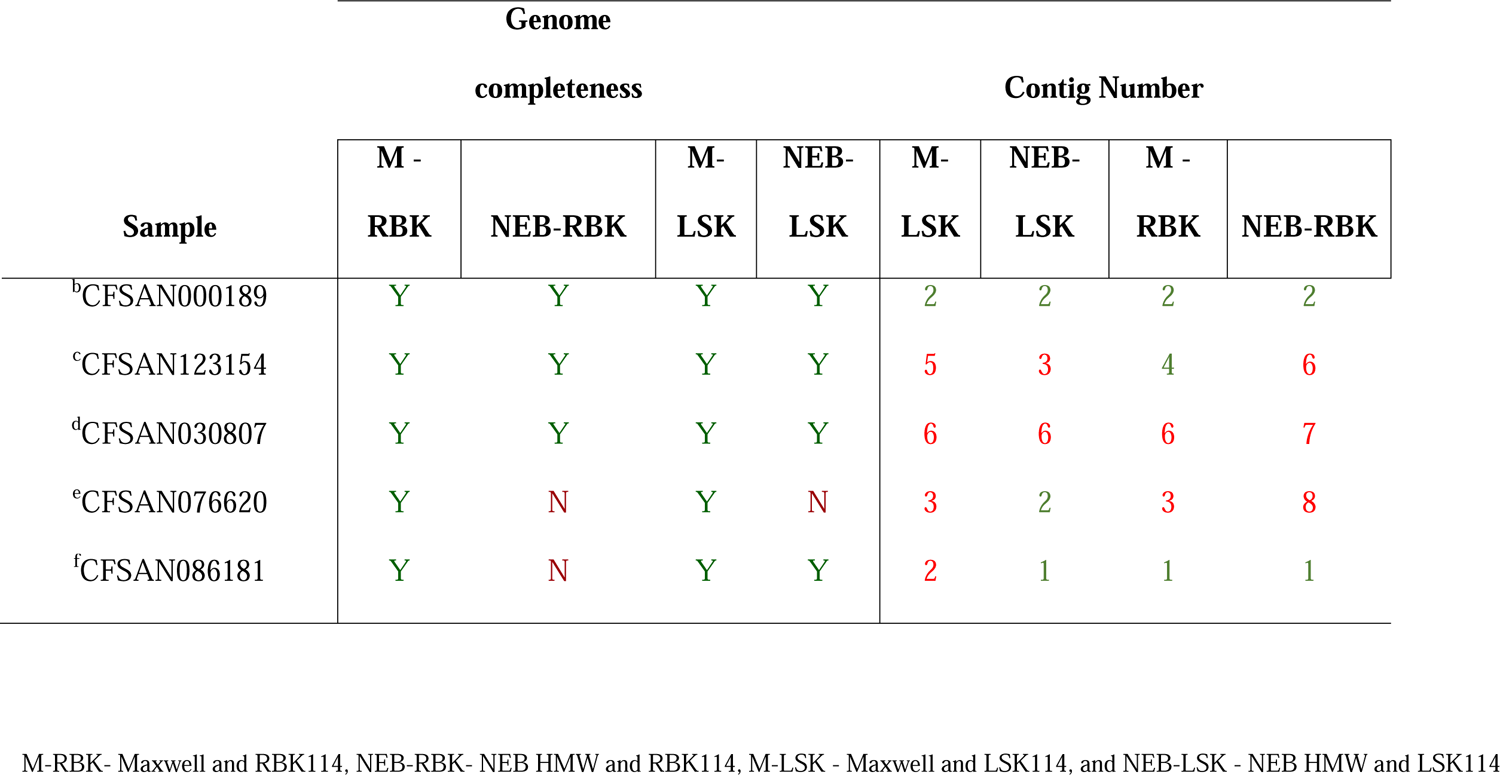
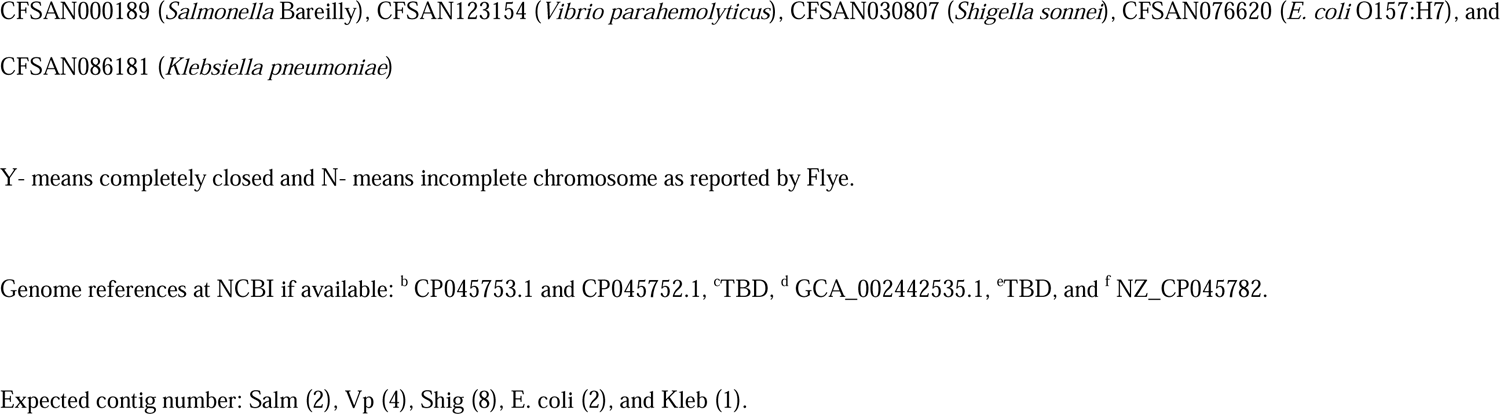
Assembly statistics showing genome completeness and contig number for each sample sequenced using any of the 4 combinations tested.

**Table 5.** Percentage of ORF genes recovered using any of the testing combinations for each strain per run. Indels and missing genes are also shown.

| % ORF genes recovered intact using wgMLST scheme<br>(genes missing/indels) |  |  |  |  |  |
| --- | --- | --- | --- | --- | --- |
|  | M-LSK | M-RBK | NEB-LSK | NEB-RBK | total |
| Samples | FAW43075 | FAW25027 | FAW42567 | FAW18242 | genes<sup>a</sup> |
| CFSAN000189 | 99.2 (1/30) | 99.1 (2/37) | 99.1 (1/36) | 99.3 (0/30) | 4296 |
| CFSAN123154 | 99.5 (2/52) | 99.2 (5/57) | 99.5 (9/44) | 98.9 (2/79) | 4440 |
| CFSAN030807 | 99.8 (2/8) | 100 (1/0) | 99.8 (1/6) | 99.7 (2/11) | 3669 |
| CFSAN076620 | 99.7 (1/34) | 99.6 (4/35) | 99.8 (1/31) | 99.5 (8/66) | 4639 |
| CFSAN086181 | 99.6 (2/21) | 99.8 (0/10) | 99.7 (1/15) | 99.7 (4/20) | 4841 |

<sup>a</sup>Total genes in each wgMLST scheme (reference genome specific) used for the analysis.

**Table 6.** SNP matrix of the second wgMLST analysis for CFSAN000189, showing SNP differences among the different combinations of ONT sequencing and the PacBio and Hybrid assemblies.

|  | 1 | 2 | 3 | 4 | 5 | 6 | 7 | 8 | 9 |
| --- | --- | --- | --- | --- | --- | --- | --- | --- | --- |
| 1 CFSAN000189_FAW25027_f | 0 | 119 | 133 | 131 | 100 | 100 | 100 | 100 | 100 |
| 2 CFSAN000189_FAW42567_f | 119 | 0 | 143 | 137 | 109 | 109 | 109 | 109 | 109 |
| 3 CFSAN000189_FAW43075_f | 133 | 143 | 0 | 126 | 124 | 124 | 124 | 124 | 124 |
| 4 CFSAN000189_FAW18242_f | 131 | 137 | 126 | 0 | 117 | 117 | 117 | 117 | 117 |
| 5 CFSAN000189_CP045753 | 100 | 109 | 124 | 117 | 0 | 0 | 0 | 0 | 0 |
| 6 CFSAN000189_M1 | 100 | 109 | 124 | 117 | 0 | 0 | 0 | 0 | 0 |
| 7 CFSAN000189_M2 | 100 | 109 | 124 | 117 | 0 | 0 | 0 | 0 | 0 |
| 8 CFSAN000189_M3 | 100 | 109 | 124 | 117 | 0 | 0 | 0 | 0 | 0 |
| 9 CFSAN000189_NCBI | 100 | 109 | 124 | 117 | 0 | 0 | 0 | 0 | 0 |

NCBI – PacBio, CP045753 – Hybrid ONT +Illumina, M- MiSeq.

### In silico analyses

The data generated from the 4 runs was compared for diverse in silico tests or assays: a) correct species identification (accuracy, specificity, and sensitivity), b) genotyping methods (serotyping, AMR, virulence, toxin, MLST), and c) phylogenomic clustering methods measuring the specificity and sensitivity for cluster assignment.

a. **correct species identification (accuracy, specificity, and sensitivity).** We used Kraken 2 implemented in the Galaxy Trakr website [19] for species identification. The assemblies for each strain from each run was ran through it and each species was identified correctly (100% accuracy, specificity, and sensitivity).
b. **genotyping methods (serotyping, AMR, virulence, toxin, MLST).** Regarding the serotyping, only two serotyping schemes were available for the 5 different species included in this pilot study (*Salmonella* and *E. coli*). There was a 100% agreement with the serotype predictions as all *Salmonella* and *E. coli* ONT assemblies were identified as serotype Bareilly and O157:H7, respectively. Similarly, *in silico* sequence type (ST) determination yielded identical results for ONT assembly (ST-909 for CFSAN000189, ST-635 for CFSAN123154, ST-152 for CFSAN030807, ST-11 for CFSAN076620, and ST-629 for CFSAN086181), except for one assembly (CFSAN000189) that showed an incomplete MLST profile due to an indel in the *sucA* gene (NEB-LSK114). All ONT assemblies for each strain agreed 100% for the presence of specific antimicrobial resistance (AMR) genes compared to the reference genome for each strain. CFSAN000189 carried a single copy of the *aac(6’)-Iaa* gene, CFSAN123154 carried a single AMR gene (*blaCARB-*36), CFSAN076620 carried 2 copies of the *catA1* gene, and CFSAN086181 carried 5 genes (*blaSHV-110, blaSHV-81, fosA6, OqxB* and *OqxA*). An Issue arose with the copy number for the AMR genes for strain CFSAN030807, which carried four AMR genes [aph(6)-Id, aph(3’’)-Ib, sul2, and tet(A)]. However, in the assemblies for runs M-LSK114 and M-RBK114, the plasmid carrying the AMR genes was double in size compared to the original plasmid, likely due to an assembly issue (we have observed this same issue in many Q20+ ONT assemblies), resulting in the same sequence being concatenated twice. For this pilot study, *in silico* analyses were conducted on ONT assemblies of the five strains to determine the presence of virulence and toxin genes. The genes reported for each species in the virulence factors database (VFDB) task template included in the Ridom Seqsphere software were utilized for these analyses, except for *E. coli*, where Virulence Finder at DTU was used instead. The reference genome of *Salmonella enterica* subsp. *enterica* (CFSAN000189) Bareilly contained 105 virulence genes, all of which were consistently identified in ONT assemblies. Similarly, the *Vibrio parahemolyticus* (CFSAN123154) strain, characterized as *tdh*-/*trh*+, contained 42 virulence genes in its reference genome, all of which were also found in every ONT assembly. Additionally, the *Shigella sonnei* (CFSAN030807) strain’s reference genome contained 10 virulence genes, all of which were consistently identified in ONT assemblies. This strain has lost a larger plasmid that carried 42 additional virulence genes compared to the current genome available at NCBI. In the case of *E. coli* (CFSAN076620), the reference genome contained 20 relevant virulence genes, which were consistently detected across different ONT combinations, with only one gene (*toxB*) missing in certain ONT assemblies, specifically NEB-RBK114 and NEB-LSK114. Finally, the reference genome of *Klebsiella pneumoniae* (CFSAN086181) harbored 52 virulence genes, all of which were present in every ONT assembly.
c. **phylogenomic clustering methods measuring the specificity and sensitivity for cluster assignment.** For this last step we used a wgMLST approach (using as reference genome the known genome sequence of the tested strains) to determine specificity and sensitivity for calling variants (SNPS and/or indels), accuracy of entire tree OR specific splits (e.g., leading to an outbreak lineage). The entire workflow is depicted in figure 1.

### wgMLST analyses for CFSAN000189 (*Salmonella* Bareilly)

We conducted two different wgMLST analyses: 1) against a collection of closely related *Salmonella* Bareilly genomes, and 2) against the reference genome for that same strain (obtained by PacBio or by a hybrid assembly of ONT and Illumina data). Total genes used in the wgMLST were 4312 loci. Of those, 3991 loci (93%) were identical among all samples (including the PacBio and hybrid genome). The ONT assemblies obtained with the Q20+ chemistry (V14) by either combination clustered with the corresponding genome in a SNP NJ tree after a wgMLST analysis (Figure 2A), albeit with at least 100 to 124 SNPs and 30-39 indels, compared to the reference genome (Figure 2B and Table 4). The SNP differences matrix is shown in Table 4. We must add that this strain was sequenced using the same initial bacterial culture, thus the DNA must be the same in terms of methylations. The complete NJ SNP tree can be found on supplementary figure 1. Additionally, the Illumina sequences [obtained using a MiSeq (M)] assembled were also included in the wgMLST analysis and were indistinguishable from the PacBio or ONT hybrid assemblies.

**Figure 2.**
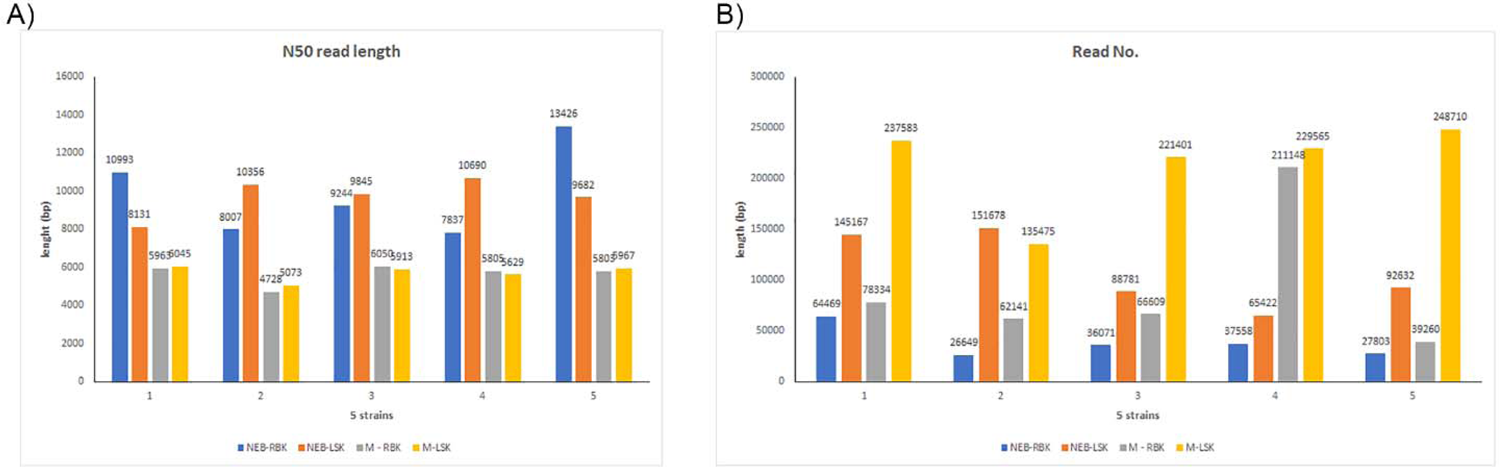
Statistics regarding N50 Length and read numbers per run for the initial testing.

### wgMLST analyses for CFSAN123154 (*Vibrio parahemolyticus*)

We conducted the same two different wgMLST analyses for the *Vibrio parahemolyticus* strain as described for *Salmonella:* 1) against a collection of closely related *Vibrio parahaemolyticus* genomes, and 2) against the reference genome for that same strain (obtained by PacBio). Total genes used in the wgMLST were 4440 loci. Of those, 4078 loci (92%) were identical among all samples (same DNA prepared with different library kit and ran in different flow cells). The ONT assemblies obtained with the Q20+ chemistry (V14) by either combination clustered with the corresponding genome in a SNP NJ tree after a wgMLST analysis (Figure 3A), albeit with at least 100 to 133 SNPs and 53 to 81 indels, compared to the reference genome (Figure 3B and Table 7). The complete NJ SNP tree can be found on supplementary figure 2. The SNP differences matrix is shown in Table 7.

**Figure 3.**
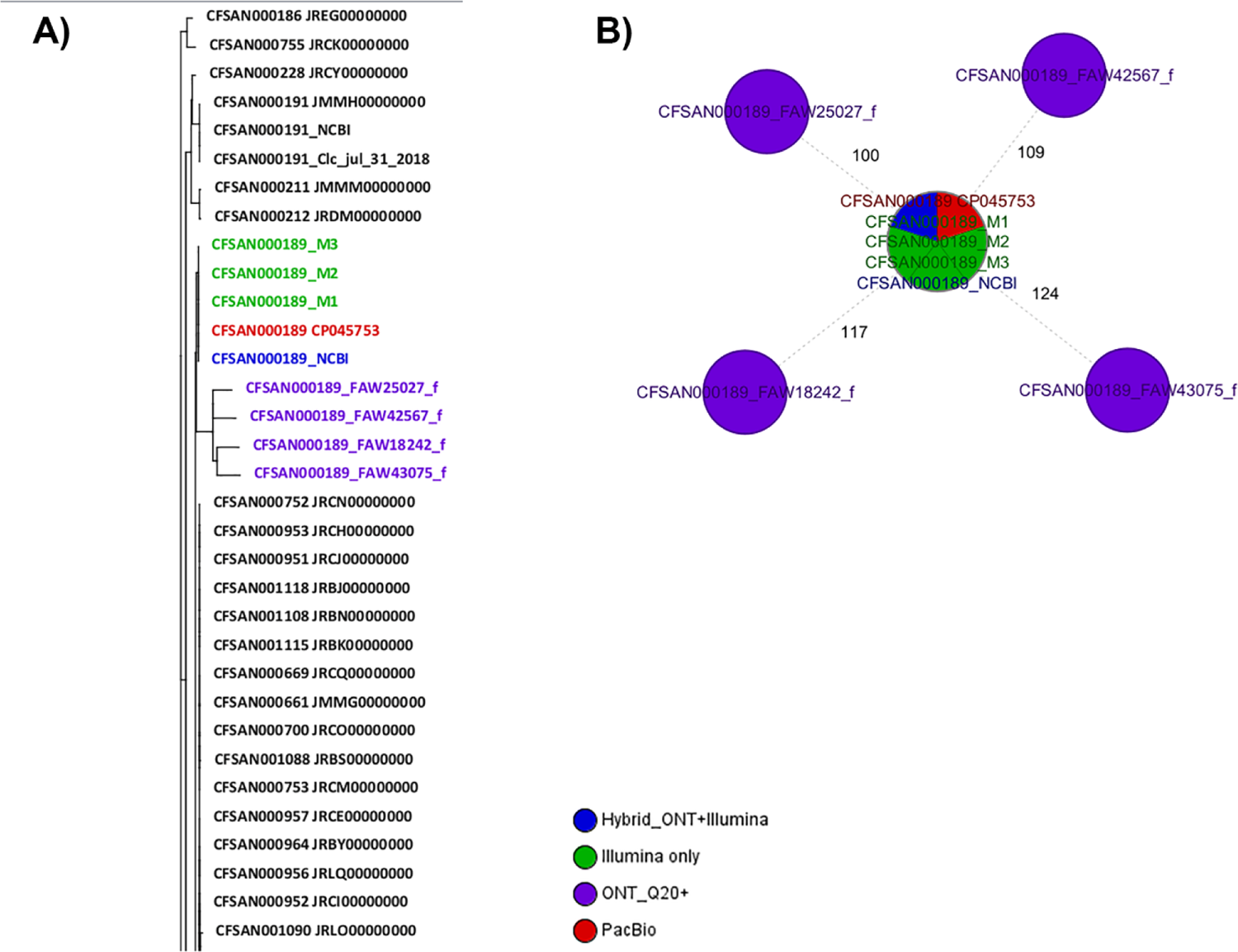
Results of the wgMLST analyses for CFSAN000189. A) Snapshot of the NJ tree generated from the wgMLST analysis of the ONT assemblies obtained by the different combination tested against a set of known genomes closely related to that same strain (ST909). B) Minimum spanning tree (MST) showing the differences between the different CFSAN000189 assemblies obtained by different sequencing technologies. The complete NJ tree can be found in supplementary figure 1.

**Figure 4.**
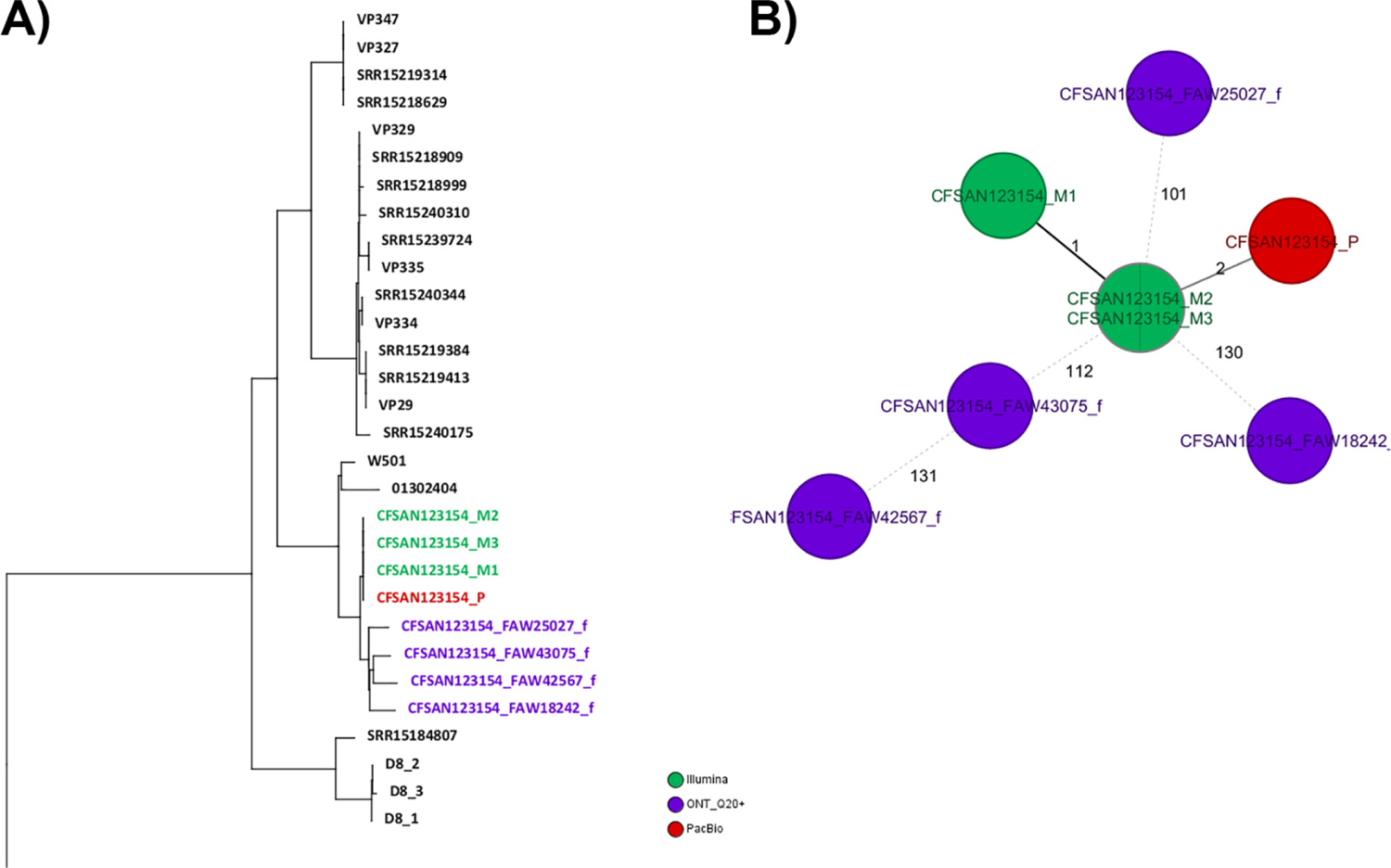
Results of the wgMLST analyses for CFSAN123154. A) NJ tree generated from the wgMLST analysis of the ONT assemblies obtained by the different combination tested against a set of known genomes closely related to that same strain. B) Minimum spanning tree (MST) showing the differences between the different CFSAN123154 assemblies obtained by different sequencing technologies. The complete NJ tree can be found in supplementary figure 2.

**Table 7.** SNP matrix of the second wgMLST analysis for CFSAN123154, showing SNP differences among the different combinations of ONT sequencing and the PacBio assembly.

|  | 1 | 2 | 3 | 4 | 5 | 6 | 7 | 8 |
| --- | --- | --- | --- | --- | --- | --- | --- | --- |
| 1 CFSAN123154_FAW25027_f | 0 | 159 | 101 | 102 | 101 | 101 | 138 | 153 |
| 2 CFSAN123154_FAW18242_f | 159 | 0 | 130 | 131 | 130 | 130 | 160 | 171 |
| 3 CFSAN123154_M2 | 101 | 130 | 0 | 1 | 0 | 2 | 112 | 136 |
| 4 CFSAN123154_M1 | 102 | 131 | 1 | 0 | 1 | 3 | 113 | 137 |
| 5 CFSAN123154_M3 | 101 | 130 | 0 | 1 | 0 | 2 | 112 | 136 |
| 6 CFSAN123154_P | 101 | 130 | 2 | 3 | 2 | 0 | 112 | 136 |
| 7 CFSAN123154_FAW43075_f | 138 | 160 | 112 | 113 | 112 | 112 | 0 | 131 |
| 8 CFSAN123154_FAW42567_f | 153 | 171 | 136 | 137 | 136 | 136 | 131 | 0 |

P – PacBio, M- MiSeq.

### wgMLST analyses for CFSAN030807 (*Shigella sonnei*)

We conducted two different wgMLST analyses for the *Shigella sonnei* strain as described for *Salmonella*, 1) against a collection of closely related *Shigella sonnei* genomes (ST-152), and 2) against the reference genome for that same strain (obtained by PacBio). Total genes used in the wgMLST were 3699 loci. Of those, 3657 loci (99%) were identical among all samples (same DNA prepared with different library kit and ran in different flow cells). The ONT assemblies obtained with the Q20+ chemistry (V14) by either combination clustered with the corresponding genome in a SNP NJ tree after a wgMLST analysis (Figure 5A), with at least 0 to 2 SNPs and 1 to 13 indels, compared to the reference genome (Figure 5B and Table 8). The SNP differences matrix is shown in Table 8. We must add that this strain was sequenced using the same initial bacterial culture, so the DNA must be the same in terms of methylations. The complete NJ SNP tree can be found in supplementary figure 3.

**Figure 5.**
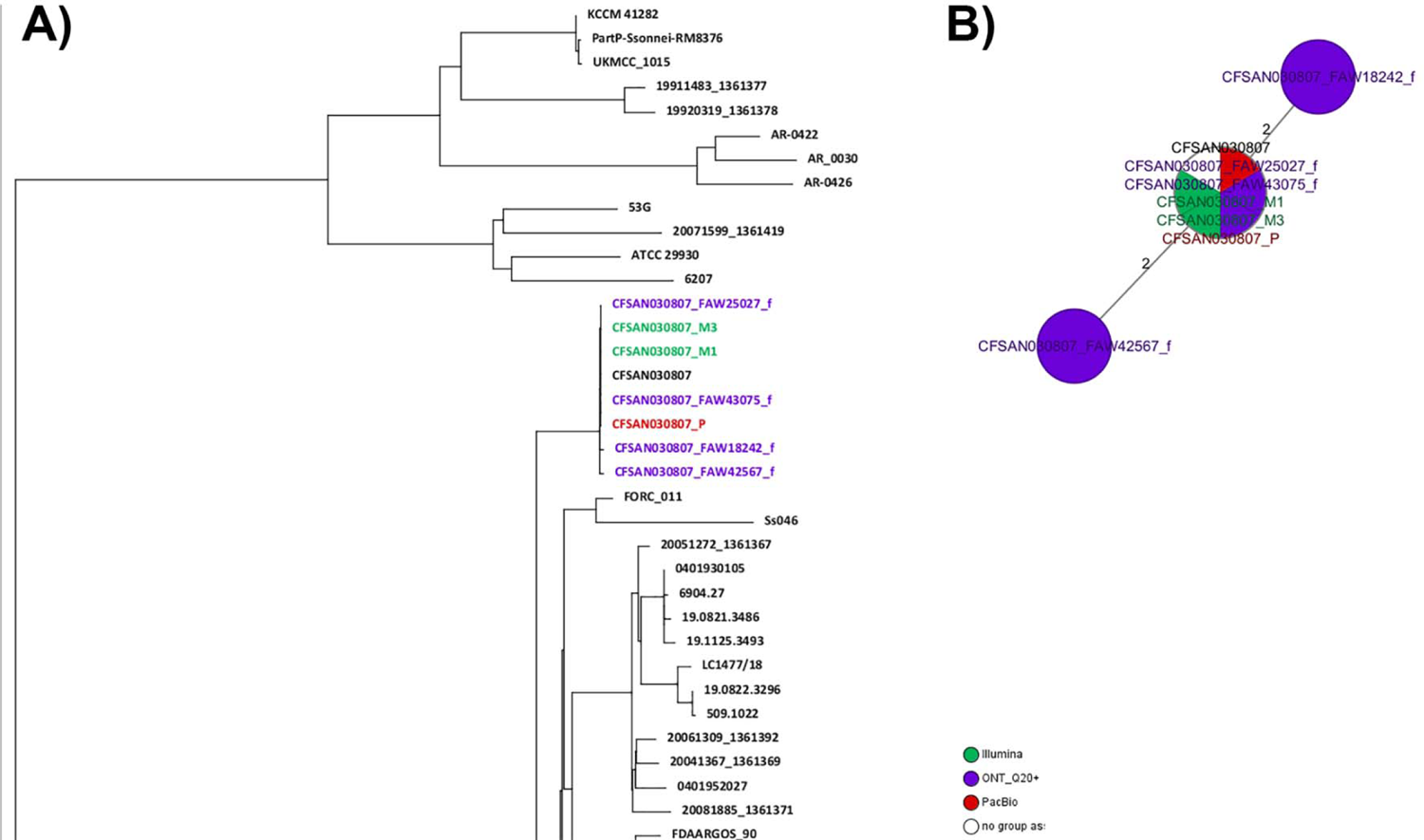
Results of the wgMLST analyses for CFSAN030807. A) Snapshot of the NJ tree generated from the wgMLST analysis of the ONT assemblies obtained by the different combination tested against a set of known genomes (156) closely related to that same strain (ST152). B) Minimum spanning tree (MST) showing the differences between the different CFSAN030807 assemblies obtained by different sequencing technologies. The complete NJ tree can be found in supplementary figure 3.

**Table 8.**
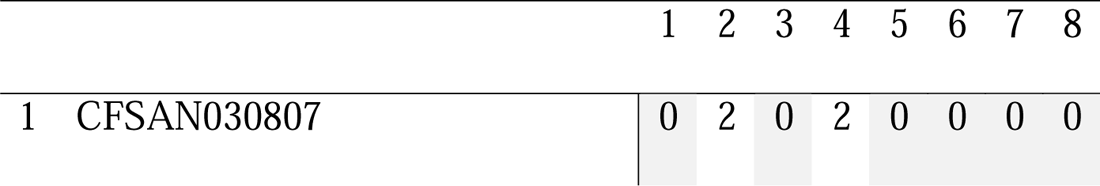
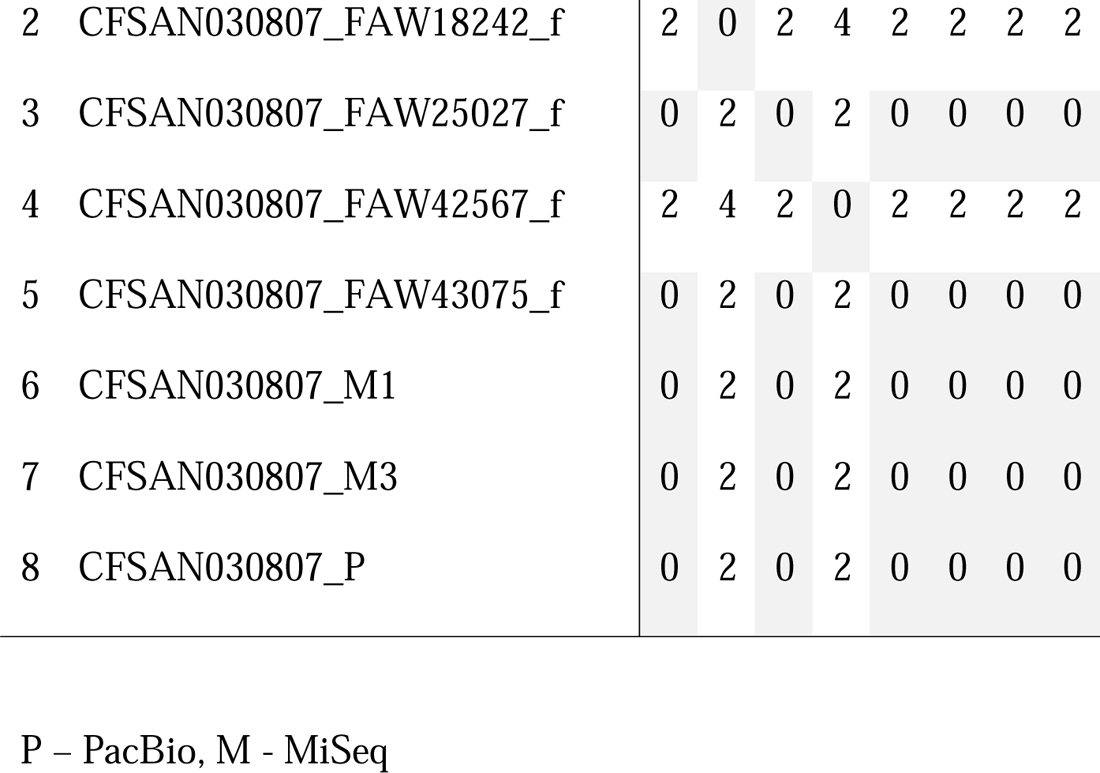
SNP matrix of the second wgMLST analysis for CFSAN030807, showing SNP differences among the different combinations of ONT sequencing and the PacBio assembly.

|  | 1 | 2 | 3 | 4 | 5 | 6 | 7 | 8 |
| --- | --- | --- | --- | --- | --- | --- | --- | --- |
| 1 CFSAN030807 | 0 | 2 | 0 | 2 | 0 | 0 | 0 | 0 |

|  |  |  |  |  |  |  |  |  |  |
| --- | --- | --- | --- | --- | --- | --- | --- | --- | --- |
| 2 | CFSAN030807_FAW18242_f | 2 | 0 | 2 | 4 | 2 | 2 | 2 | 2 |
| 3 | CFSAN030807_FAW25027_f | 0 | 2 | 0 | 2 | 0 | 0 | 0 | 0 |
| 4 | CFSAN030807_FAW42567_f | 2 | 4 | 2 | 0 | 2 | 2 | 2 | 2 |
| 5 | CFSAN030807_FAW43075_f | 0 | 2 | 0 | 2 | 0 | 0 | 0 | 0 |
| 6 | CFSAN030807_M1 | 0 | 2 | 0 | 2 | 0 | 0 | 0 | 0 |
| 7 | CFSAN030807_M3 | 0 | 2 | 0 | 2 | 0 | 0 | 0 | 0 |
| 8 | CFSAN030807_P | 0 | 2 | 0 | 2 | 0 | 0 | 0 | 0 |

P – PacBio, M - MiSeq

### wgMLST analyses for CFSAN076620 (*E. coli* O157:H7)

We conducted two different wgMLST analyses for the *E. coli* strain similar as described for *Salmonella*, 1) against a collection of closely related *E. coli O157:H7* genomes (ST-11), and 2) against the reference genome for that same strain (obtained by PacBio). Total genes used in the wgMLST were 4639 loci. Of those, 4500 loci (97%) were identical among all samples (same DNA prepared with different library kit and ran in different flow cells). The ONT assemblies obtained with the Q20+ chemistry (V14) by either combination clustered with the corresponding genome in a SNP NJ tree after a wgMLST analysis (Figure 6A), albeit with at least 3 to 28 SNPs and 32 to 74 indels, compared to the reference genome (Figure 6B and Table 9). The SNP differences matrix is shown in Table 9. We must add that this strain was sequenced using the same initial bacterial culture, so the DNA must be the same in terms of methylations.

**Figure 6.**
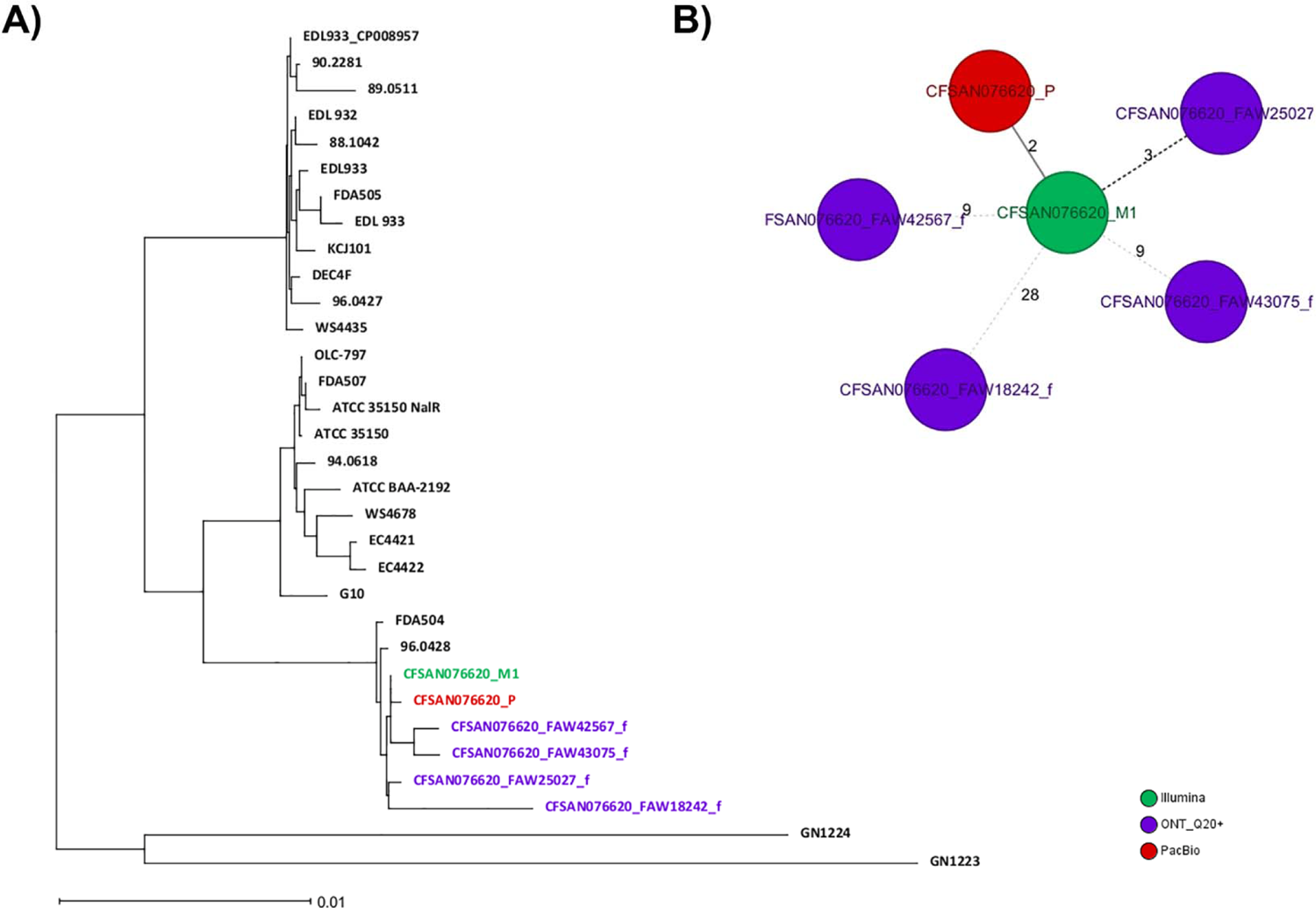
Results of the wgMLST analyses for CFSAN076620. A) NJ tree generated from the wgMLST analysis of the ONT assemblies obtained by the different combination tested against a set of known genomes (54) closely related to that same strain (ST629). B) Minimum spanning tree (MST) showing the differences between the different CFSAN076620 assemblies obtained by different sequencing technologies.

**Table 9.** SNP matrix of the second wgMLST analysis for CFSAN076620, showing SNP differences among the different combinations of ONT sequencing and the PacBio assembly.

|  | 1 | 2 | 3 | 4 | 5 | 6 |
| --- | --- | --- | --- | --- | --- | --- |
| 1 CFSAN076620_FAW18242_f | 0 | 28 | 36 | 28 | 30 | 35 |
| 2 CFSAN076620_M1 | 28 | 0 | 9 | 3 | 2 | 9 |
| 3 CFSAN076620_FAW42567_f | 36 | 9 | 0 | 12 | 11 | 9 |
| 4 CFSAN076620_FAW25027_f | 28 | 3 | 12 | 0 | 5 | 10 |
| 5 CFSAN076620_P | 30 | 2 | 11 | 5 | 0 | 11 |
| 6 CFSAN076620_FAW43075_f | 35 | 9 | 9 | 10 | 11 | 0 |

P – PacBio, M - MiSeq

### wgMLST analyses for CFSAN086181 (*Klebsiella pneumoniae*)

We conducted two different wgMLST analyses for the *Klebsiella pneumoniae* strain similar as described for *Salmonella*, 1) against a collection of closely related *Klebsiella pneumoniae* genomes, and 2) against the reference genome for that same strain (obtained by PacBio). Total genes used in the wgMLST were 4841 loci. Of those, 4778 loci (98.7%) were identical among all samples (same DNA prepared with different library kit and ran in different flow cells). The ONT assemblies obtained with the Q20+ chemistry (V14) by either combination clustered with the corresponding genome in a SNP NJ tree after a wgMLST analysis (Figure 7A), with 1 to 9 SNPs and 10 to 24 indels compared to the reference genome (Figure 7B and Table 4). The SNP differences matrix is shown in Table 10. We must add that this strain was sequenced using the same initial bacterial culture, so the DNA must be the same in terms of methylations. The complete NJ SNP tree can be found in supplementary figure 4.

**Figure 7.**
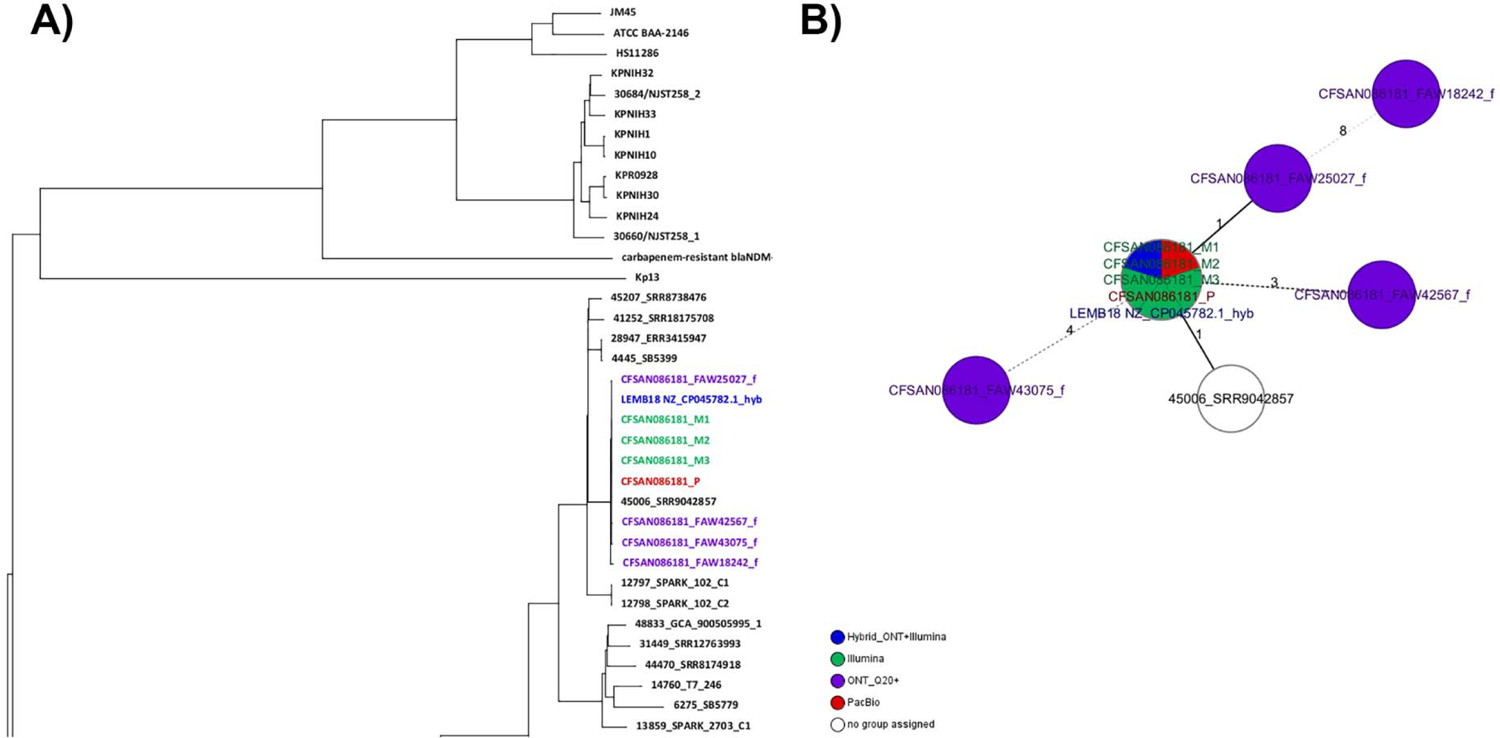
Results of the wgMLST analyses for CFSAN086181. A) Snapshot of the NJ tree generated from the wgMLST analysis of the ONT assemblies obtained by the different combination tested against a set of known genomes (54) closely related to that same strain (ST629). B) Minimum spanning tree (MST) showing the differences between the different CFSAN086181 assemblies obtained by different sequencing technologies. The complete NJ tree can be found in supplementary figure 4.

**Table 10.**
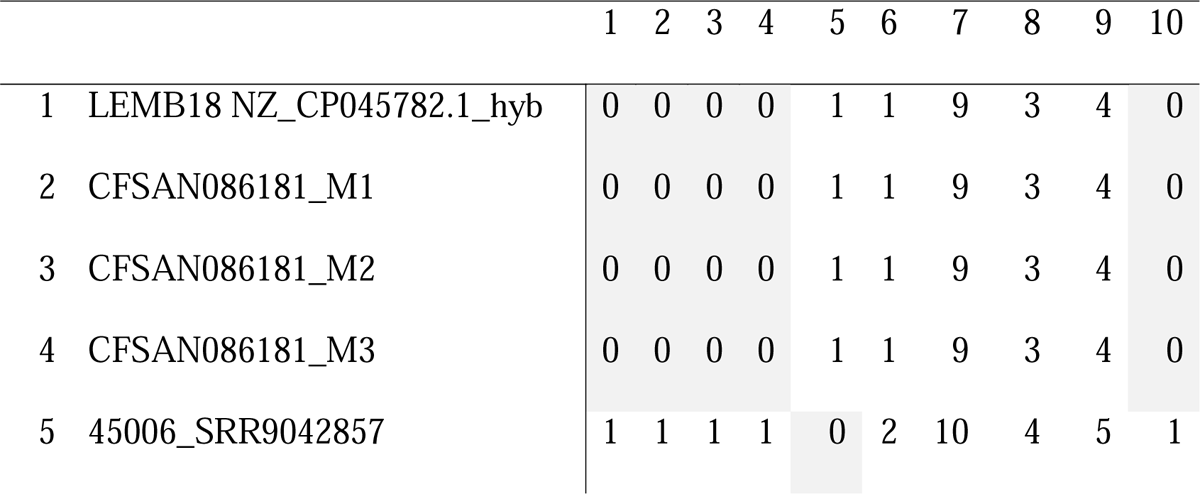
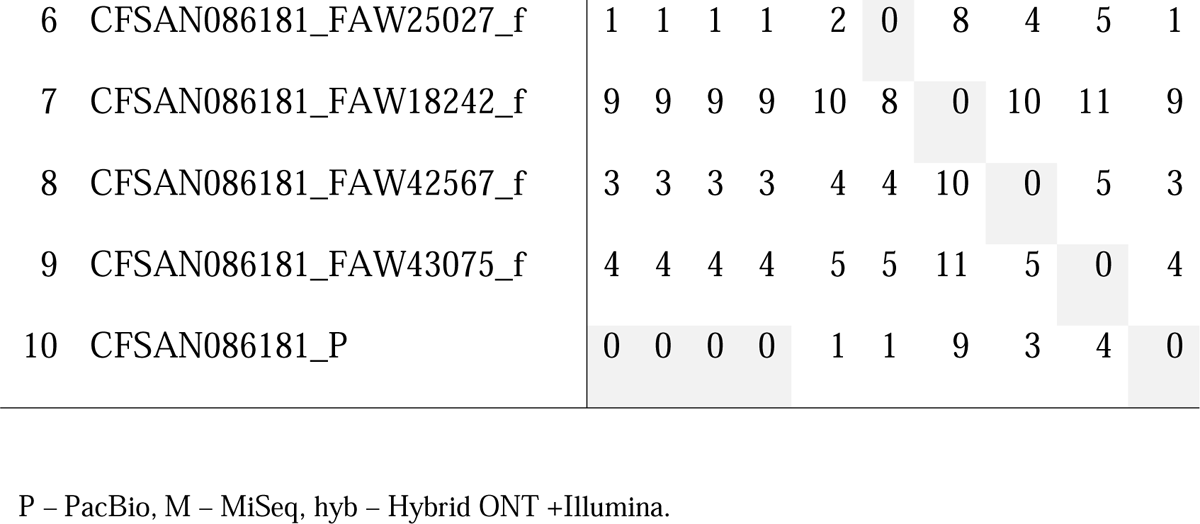
SNP matrix of the second wgMLST analysis for CFSAN086181, showing SNP differences among the different combinations of ONT sequencing and the PacBio, Hybrid, and Illumina assemblies.

|  | 1 | 2 | 3 | 4 | 5 | 6 | 7 | 8 | 9 | 10 |
| --- | --- | --- | --- | --- | --- | --- | --- | --- | --- | --- |
| 1 LEMB18 NZ_CP045782.1_hyb | 0 | 0 | 0 | 0 | 1 | 1 | 9 | 3 | 4 | 0 |
| 2 CFSAN086181_M1 | 0 | 0 | 0 | 0 | 1 | 1 | 9 | 3 | 4 | 0 |
| 3 CFSAN086181_M2 | 0 | 0 | 0 | 0 | 1 | 1 | 9 | 3 | 4 | 0 |
| 4 CFSAN086181_M3 | 0 | 0 | 0 | 0 | 1 | 1 | 9 | 3 | 4 | 0 |
| 5 45006_SRR9042857 | 1 | 1 | 1 | 1 | 0 | 2 | 10 | 4 | 5 | 1 |

|  |  |  |  |  |  |  |  |  |  |  |  |
| --- | --- | --- | --- | --- | --- | --- | --- | --- | --- | --- | --- |
| 6 | CFSAN086181_FAW25027_f | 1 | 1 | 1 | 1 | 2 | 0 | 8 | 4 | 5 | 1 |
| 7 | CFSAN086181_FAW18242_f | 9 | 9 | 9 | 9 | 10 | 8 | 0 | 10 | 11 | 9 |
| 8 | CFSAN086181_FAW42567_f | 3 | 3 | 3 | 3 | 4 | 4 | 10 | 0 | 5 | 3 |
| 9 | CFSAN086181_FAW43075_f | 4 | 4 | 4 | 4 | 5 | 5 | 11 | 5 | 0 | 4 |
| 10 | CFSAN086181_P | 0 | 0 | 0 | 0 | 1 | 1 | 9 | 3 | 4 | 0 |

P – PacBio, M – MiSeq, hyb – Hybrid ONT +Illumina.

### Testing the new software update (MinKNOW 23.04.5) for same samples using the RBK-114 kit

We tested the new software update (MinKNOW 23.04.5) using the RBK-114 kit on the same samples. During our testing of the new Q20+ chemistry, ONT released an update for the MinKNOW software (v23.04.5), which incorporated a new basecall model: dna_r10.4.1_e8.2_5khz_400bps_sup@v4.2.0. ONT claimed that there were no differences in the accuracy of the SUP 400 bps compared to the previous SUP at 260 bps. We conducted tests using the same five DNA strains and the rapid barcoding kit (RBK-114). The output was very similar to the observed RBK runs (741.66k reads and 3.9 Gb, flow cell number FAW81132). The procedure for data analysis remained consistent, employing two different wgMLST analyses for each strain: 1) against a collection of closely related genomes, and 2) against the reference genome for the same strain. Every strain clustered in the same cluster for each individual species as observed with the SUP 260 bps. Regarding indels and SNPs: - CFSAN000189 had 32 SNPs and 56 indels compared to its reference genome, - CFSAN123154 had 77 SNPs and 78 indels compared to its reference genome, - CFSAN030807 had 0 SNPs and 5 indels compared to its reference genome, - CFSAN076620 had 3 SNPs and 27 indels compared to its reference genome, and - CFSAN086181 had 2 SNPs and 7 indels compared to its reference genome.

## DISCUSSION

This study was conducted to address the limitations of current FDA pathogen sequencing methods, which are time consuming and require expensive equipment. This research aimed to evaluate whether Oxford Nanopore Technologies (ONT) MinION with Q20+ chemistry can offer rapid and accurate identification of foodborne pathogens, including SNP differences, serotypes, and antimicrobial resistance genes. The goal was to assess if ONT Q20+ technology could provide near real-time pathogen identification, potentially overcoming the drawbacks of current methodologies. The study aimed to bridge advanced genomic technologies with regulatory science, enabling more effective responses to foodborne disease threats and enhancing public health outcomes.

To achieve this, the study evaluated ONT Q20+ chemistry using five well-characterized strains representing diverse foodborne pathogen traits. Various DNA extraction (Maxwell and NEB HMW) and library preparation methods (RBK-114 and LSK-114) were compared to optimize sequencing outcomes. We observed that any combination was sufficient to produced high quality bacterial genomes using the Flye assembler as observed by many other authors under similar conditions [11–13]. Variations in assembly length and genome completeness was observed across different combinations of extraction and sequencing kits. Most genomes achieved high completeness (>99% ORF recovery) with ONT Q20+ as has been observed previously [13, 14, 23], but some variations in assembly length and missing ORFs were noted, suggesting the need for further optimization. We also observed duplication of small plasmids in some cases when using the Flye assembler as observed by many others [12, 24–26]. Specifically for strain CFSAN030807, where the expected small plasmid of 8,401 bp carrying four AMR genes was 16,802 bp in size for the assemblies of M - RBK114 and M - LSK114. So, the reason the plasmid is double in size seems random or perhaps the DNA extracted with that specific kit created smaller fragments for that plasmid and were randomly miss-assembled due to some minor SNP differences.

Regarding the *in silico* analyses, the Q20+ nanopore sequencing chemistry demonstrated 100% for accurate species identification, genotyping (including serotyping and AMR gene detection), and virulence factor identification comparable to Illumina sequencing. Phylogenomic clustering methods (wgMLST) showed that ONT assemblies clustered with reference genomes, although some indels and SNP differences were observed with some bacterial species and less pronounced with others. Supplementing with Illumina data (results not shown) helped resolve these differences, highlighting the need for accurate variant calling. Other authors have similar results for other enterobacterial species (*Citrobacter freundii, Klebsiella pneumoniae, Enterobacter hormaechei, Enterobacter cloacae,* and *Escherichia coli*) and noticed that there was still a need to supplement with Illumina data in order to correct those indels or SNPs [12]. Despite these variations, the overall clustering and similarity to reference genomes highlight the robustness of nanopore sequencing results for outbreak investigations. The SNPs observed were randomly distributed in the genome of some of the analyzed strains. This addition clarifies that any observed SNP differences in the phylogenetic analyses likely stemmed from sequencing and analysis processes rather than genetic variations in the sampled bacteria. It’s important to note that these analyses were conducted using DNA from the same initial bacterial culture and the impact of methylations on the basecallers should have been the same for each strain and should not reflect on the SNPs observed for each run for that specific strain (supplementary figure 5).

The assembled genomes from each combination were compared against their reference genome annotated coding sequences (CDS) using a highly conservative requirement for a 100 % match, as described by Sanderson et. al (2024) [13]. Both indels and SNP variants were identified using this process. Regulatory actions demand a high level of certainty to identify an outbreak strain. We foresee the use of ONT for outbreak strain confirmation (identical – zero SNPs or highly identical less than five SNPs), and therefore require almost complete matching or no more than one or two SNPs differences when comparing to the reference genome. Even though we performed a single run per combination (except for the Maxell RBK-114 combination) we believe that it is equivalent to multiple replicates of the same sample. Because we used the same bacterial culture and the same chemistry, albeit with different library preparation kits and procedures, the final quality of the sequence data shouldn’t be affected. As mentioned by Sanderson et al 2024 [13], the output of read numbers and yield can also be affected by the number of pores, skill of the operator and library type. Furthermore, we only analyzed five strains from different species but, as mentioned by Sanderson et al. 2024, these findings may not be generalized across strains or species, as our results showed that depending on the species/strain the assemblies were more or less accurate compared to the reference (Tables 6-10).

During our initial evaluation of the Q20+ chemistry, the study encountered a major update of the basecaller. Using the RBK-114 kit, we processed the same five DNA strains with both the new and old basecall models. ONT’s claimed that there would be no difference in accuracy between the SUP 400 bps and the previous SUP 260 bps models and this was largely supported by our findings. Furthermore, our downstream analyses of the output data showed that the clustering for all strains were consistent with those obtained using the SUP 260 bps model. This showed that the new basecaller model did not affect the overall clustering and comparative accuracy in detecting SNPs and indels of the strains. Overall, our testing confirms that the update to MinKNOW 23.04.5 and its new basecall model does not compromise accuracy and performs comparably to the previous version, validating ONT’s claims. This indicates that users can confidently transition to the new software update without concerns about a loss in data quality. However, we must caution that ONT has changed their basecaller several times since the start of our evaluation and these constant changes could create a negative concern for current and potential new users.

## Conclusions

The evaluation of the Q20+ nanopore sequencing chemistry in this single laboratory evaluation study demonstrates its potential as an alternative to Illumina sequencing for rapid and comprehensive bacterial genome analysis in outbreak investigations. Standardizing the extraction and sequencing protocols is crucial for optimizing workflow and ensuring reproducibility and accuracy across different strains and outbreak scenarios. While the study encountered challenges in achieving consistent and reliable results with ONT Q20+ chemistry, the findings provide valuable insights into the technology’s potential for enhancing foodborne pathogen detection and surveillance. Continued research and optimization efforts are necessary to address the observed limitations and fully realize the impact of nanopore sequencing on public health and regulatory science. Further verification studies and optimizations are essential to integrate nanopore sequencing into routine outbreak surveillance and response protocols.

Overcoming current limitations will be crucial for incorporating these technologies into routine foodborne pathogen surveillance, ultimately leading to improved public health outcomes and more agile responses to foodborne disease threats. The accuracy of SNPs in the assemblies compared to Illumina or PacBio assemblies remains an issue. This is being addressed in future iterations of the ONT basecaller, but the improvements need to be properly assessed and verified.

## Acknowledgements.

This research was funded by the Food and Drug Administration Foods Program Intramural Funds.

## Supplementary Material

**Supplementary Figure 1.** Complete NJ tree of the wgMLST analyses for CFSAN000189.

**Supplementary Figure 2.** Complete NJ tree of the wgMLST analyses for CFSAN123154.

**Supplementary Figure 3.** Complete NJ tree of the wgMLST analyses for CFSAN030807.

**Supplementary Figure 4.**
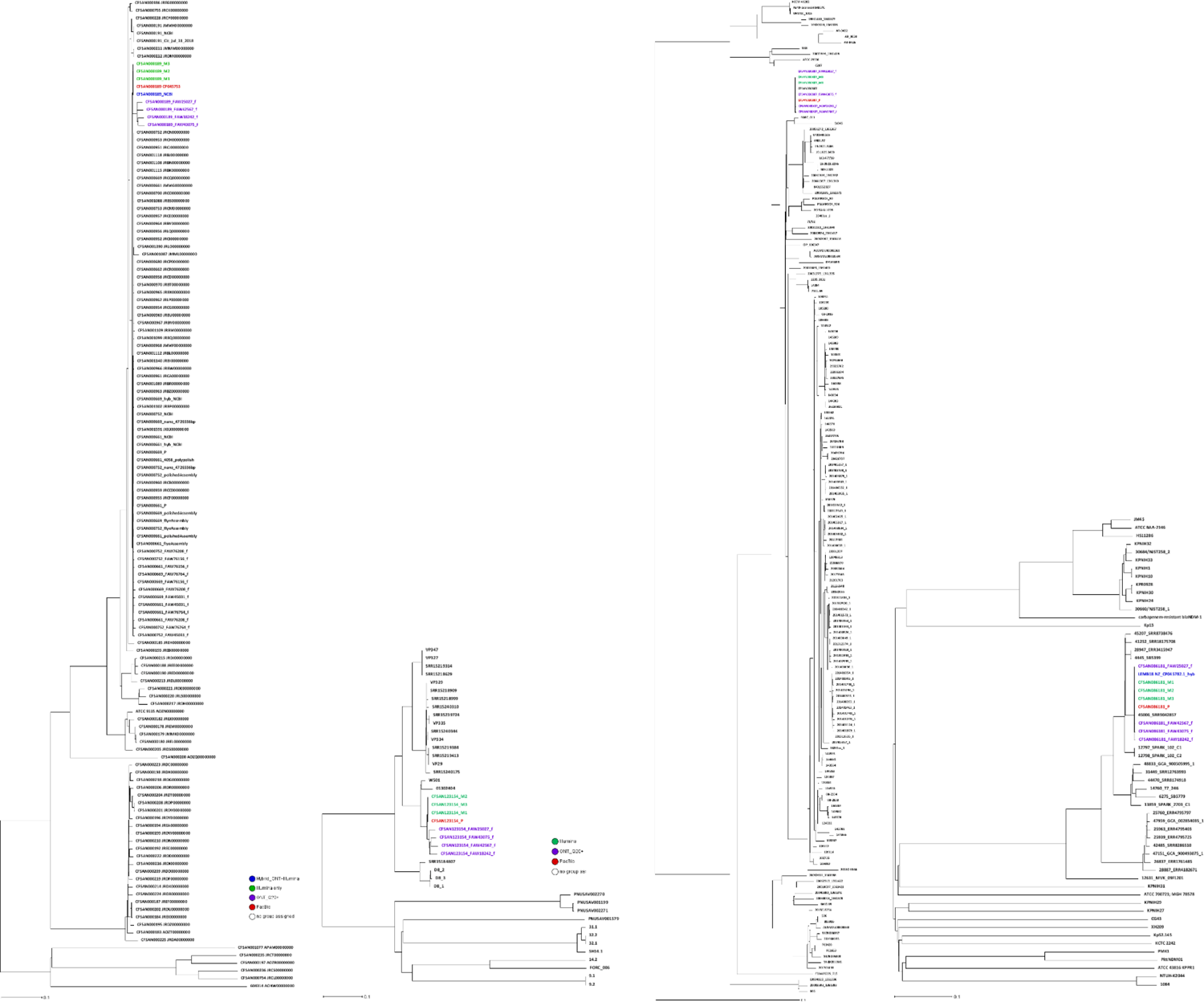
Complete NJ tree of the wgMLST analyses for CFSAN086181.

**Supplementary Figure 5.**
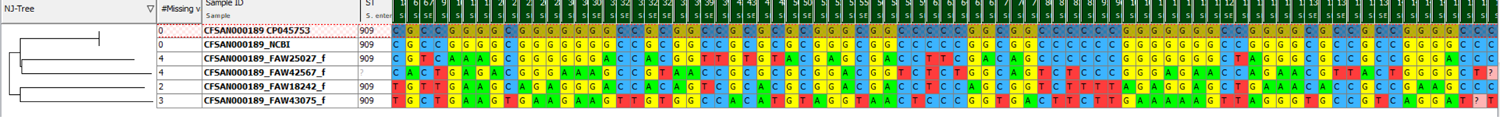
Example of the SNP differences for the same DNA ran in the four ONT library/DNA extraction combination in the initial testing for the ONT Q20+, showing that the SNPs were in different genes and positions in the genomes for the same strain.

