## Supplementary material for "Single laboratory evaluation of the (Q20+) nanopore sequencing kit for bacterial outbreak investigations"

### Supplementary Figures

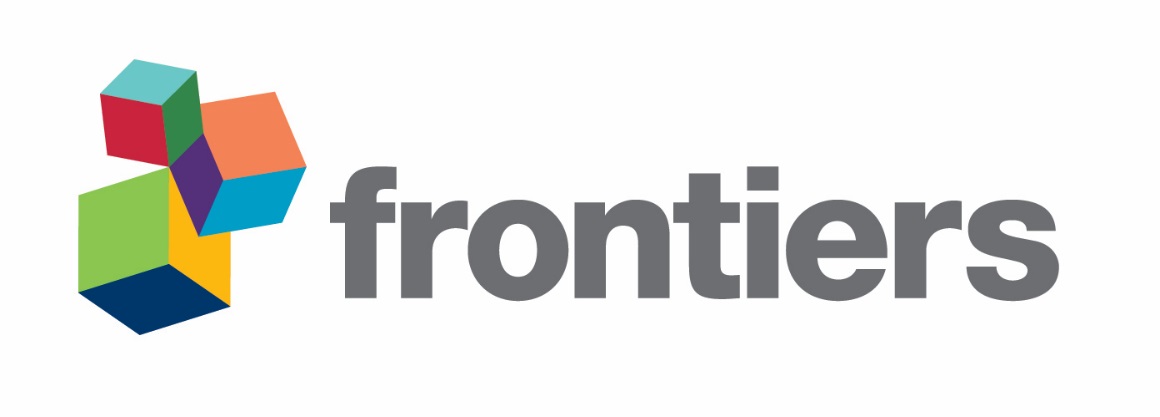

**Supplementary Figure 1**. Complete NJ tree of the wgMLST analyses for CFSAN000189**.**

**Supplementary Figure 2**. Complete NJ tree of the wgMLST analyses for CFSAN123154**.**

**Supplementary Figure 3**. Complete NJ tree of the wgMLST analyses for CFSAN030807**.**

**Supplementary Figure 4**. Complete NJ tree of the wgMLST analyses for CFSAN086181**.**

**Supplementary Figure 5**. Example of the SNP differences for the same DNA ran in the four ONT library/DNA extraction combination in the initial testing for the ONT Q20+, showing that the SNPs were in different genes and positions in the genomes for the same strain.
